# Identification of co-evolving temporal networks

**DOI:** 10.1101/303974

**Authors:** Rasha Elhesha, Aisharjya Sarkar, Christina Boucher, Tamer Kahveci

## Abstract

**Motivation:** Biological networks describes the mechanisms which govern cellular functions. Temporal networks show how these networks evolve over time. Studying the temporal progression of network topologies is of utmost importance since it uncovers how a network evolves and how it resists to external stimuli and internal variations. Two temporal networks have co-evolving subnetworks if the topologies of these subnetworks remain similar to each other as the network topology evolves over a period of time. In this paper, we consider the problem of identifying co-evolving pair of temporal networks, which aim to capture the evolution of molecules and their interactions over time. Although this problem shares some characteristics of the well-known network alignment problems, it differs from existing network alignment formulations as it seeks a mapping of the two network topologies that is invariant to temporal evolution of the given networks. This is a computationally challenging problem as it requires capturing not only similar topologies between two networks but also their similar evolution patterns.

**Results:** We present an efficient algorithm, *Tempo*, for solving identifying coevolving subnetworks with two given temporal networks. We formally prove the correctness of our method. We experimentally demonstrate that Tempo scales efficiently with the size of network as well as the number of time points, and generates statistically significant alignments—even when evolution rates of given networks are high. Our results on a human aging dataset demonstrate that Tempo identifies novel genes contributing to the progression of Alzheimer’s, Huntington’s and Type II diabetes, while existing methods fail to do so.

**Availability:** Software is available online (https://www.cise.ufi.edu/∼relhesha/temporal.zip).

**Supplementary information:** Supplementary data are available at *Bioinformatics* online.

## 1 Introduction

Biological networks describe the interaction between molecules. They are frequently represented as graphs, where the nodes correspond to the molecules (e.g., proteins or genes) and the edges correspond to their interactions (Zhu *et al*., 2007). Formally, we denote a biological network as *G* = (*V, E*) where *V* and *E* represent the set of nodes and the set of edges, respectively. Analysis of these networks enable the elucidation of cellular functions (Freyre-González *et al*., 2008), the identification of variations in cancer networks (Leiserson *et al*., 2015), and the characterization of variations in drug resistance (Charlebois *et al*., 2014). Studying biological networks led to numerous computational challenges as well as methods which address these challenges. Network alignment is one of the most important of these challenges (Flannick *et al*., 2006) as it has a profound set of applications ranging from the detection of conserved motifs to the prediction of protein functions (Clemente *et al*., 2006). This problem aims to find a mapping of the nodes of two given networks in which nodes that are similar in terms of content (i.e. homology) and interaction structure (i.e. topology) are mapped to each other. Hence, we represent the alignment between two given networks *G*_1_ = (*V*_1_, *E*_1_) and *G*_2_ = (*V*_2_, *E*_2_) as a bijection function *ψ* : *V*_1_ → *V*_2_, and the score resulting from alignment *ψ* as *score*(*G*_1_, *G*_2_|*ψ*). The network alignment problem seeks the function *ψ* that maximizes this score. We note that there are various ways to calculate the scoring function.

There are two categories of network alignment problem: local and global alignment. The former problem aims to find pairs of highly-conserved sub-networks in two given networks in which a sub-network of the query network is mapped to multiple sub-networks in the target network. Global network alignment aims to maximize the similarity in the networks in which all nodes in the query network are mapped to a set of nodes in the target network. Network alignment is a challenging task as the graph and subgraph isomorphism problems which are known to be GI and NP-hard (Cook, 1971), reduce to them. In Section 2, we give a brief review of the methods addressing the global network alignment problem as the problem we consider in this paper is associated with that problem.

Biological networks have dynamic topologies (Przytycka *et al*., 2010). There are various reasons behind this dynamic behavior. For example, genetic and epigenetic mutations can alter molecular interactions (Sadikovic *et al*., 2008), and variation in gene copy number can affect the existence of interactions (De Smith *et al*., 2008). Due to this dynamic behavior, the topology of the network that models the molecular interaction evolve over time (Holme and Saramäki, 2012). Majority of the previous work on alignment of biological networks assume the network topology is static (Singh *et al*., 2007)—an assumption that ignores the history of network evolution, and may lead to biased or incorrect analysis. For example, identifying causes and consequences of the influence of external stimuli is impossible when analyzing static topologies. To address this oversight, we define a biological network using a model that accounts for the evolution of the underlying network at consecutive time points. We refer to this model as a *temporal network* (Hulovatyy *et al*., 2015). We view this model as containing a single snapshot of the network at each time point in a sequence of time points and thus, as a time series network. More formally, we denote a temporal network with *t* consecutive time points as 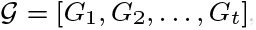, where *G_i_* = (*V, E_i_*) represents the topology of the network at the *i*th time point.

In this paper, we consider the problem of identifying *coevolving* subnetworks in a given pair of temporal networks. We say that two subnetworks are coevolving if their topologies remain similar even though their topologies evolve over time. We define this more formally as follows. We consider two input temporal networks 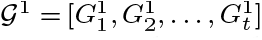 and 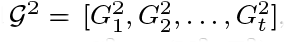, where ∀*i* ∈ {1,2, …, *t*}, 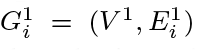 and 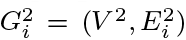 represent 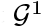 and 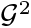 respectively at the time point *i*. Without losing generality, let 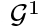 be the query (smaller) network and 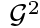 be the target network, i.e., |*V*^1^| ≤ |*V*^2^|. An alignment of 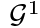 and 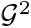 maps 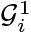 to 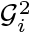 across all time points *i*. Thus, we represent the alignment of the two temporal networks 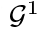 and 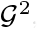 as a bijection of their nodes and denote it as a function *ψ* : *V*^1^ → *V*^2^. We compute the score of the alignment *ψ* of 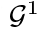 and 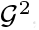, denoted with *score*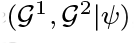, as the sum of the scores of the alignment at all time points. Hence, *score*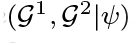 = 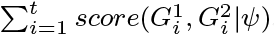. We assume 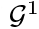 is connected at all time points, but it maybe impossible to find an alignment that is connected in the target network at all time points.

It is worth emphasizing that the temporal network alignment problem described above is dramatically different than existing network alignment problems, which can be categorized as follows: (i) pairwise alignment, (ii) multiple network alignment, and (iii) dynamic network alignment. We illustrate these problems as well as the temporal one in Figure 1. The pairwise network alignment problem (Figure 1(a)) ignores that the network topology evolves. Although the multiple alignment problem (Figure 1(b)) can consider more than two networks at once, it lacks the ability to capture the temporal changes since it treats all networks as having static topologies. The dynamic network alignment problem (Figure 1(c)) considers topological changes over time. It however, it seeks a different solution to the alignment problem at each time point. Thus, it can not identify coevolving subnetwork. A new algorithm is needed to capture such evolving characteristics. Unlike these alignment problems, temporal network alignment (Figure 1(d)) captures that network topologies coevolve over time.

**Fig. 1.**
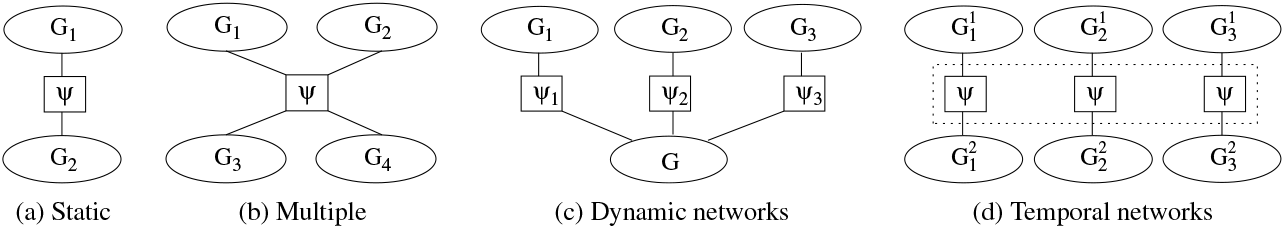
This figure represents different network alignment problems in different types of biological networks. (a) This represents the alignment between two input static networks. (b) This represents the alignment between multiple time points where each network represent a different organism. (c) This represents the alignment between two input networks where one of them is static and one of them is dynamic. Here, there exist different alignment between the static network and each version of the dynamic network. (d) This represents the alignment between two input temporal networks where each have time specific snapshots that was taken at three specific time points. Here, the alignment is persist across all time points.

### Contributions in this paper

We develop an efficient algorithm, *Tempo*, to identify coevolving subnetworks in a given pair of the temporal networks. Briefly, our algorithm first finds an initial alignment between the input networks 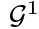 and 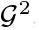 using the similarity score between pairs of aligned nodes across all time points. It then performs a dynamic programming strategy that maximizes the alignment quality (i.e. score) by repeatedly altering the alignednodesinthe targetnetwork. We demonstrate the efficiency and accuracy ofTempo using bothreal and synthetic data. We compare the running time and the quality of the alignments found by Tempo against thoseofthree existing alignment algorithms, IsoRank(Singh *et al*., 2007), MAGNA++ (Vijayan *et al*., 2015) and GHOST (Patro and Kingsford, 2012). Note that all these networks are tailored towards optimizing alignment at a single time point. To have a fair comparison, we allow each of these methods to consider each time point independently then apply the resulting alignments to all other time points and took the average. We show Tempo has competitive running time and generates significantly better alignments. We use a human brain aging (Berchtold *et al*., 2008) dataset, and integrate this dataset to analyze three phenotypes—two age related diseases (Alzheimer’s and Huntington’s) and one disease that is less prone to aging (Type II diabetes). We perform gene ontology analysis on the aligned genes reported by our algorithm and compared algorithms. Our algorithm could successfully align genes of the phenotype query (i.e. the underlying disease) to strongly related genes in the target network despite their evolving topologies unlike other algorithms. Consequently, we could predict disease-related genes based on the generated alignment using tempo which suggests that Tempo generates alignments that reflect the evolution of nodes topologies through time as well as their homological similarities while other methods only focuses on static and independent topologies. Lastly, we observe that alignments of agerelatedphenotype is significantly higher than alignment of non age phenotype which reflects their high evolution rates and shows that Tempo could identify between different queries.

## 2 Related Work

One of the key studies on pairwise global network alignment is IsoRank (Singh *et al*., 2007), which is based on the conjecture that two nodes should be matched if their respective neighbors can also be matched. It formulates the alignment as an eigenvalue problem and computes the similarity between pairs of nodes from two given networks as a combination of their homological and topological similarities. It obtains the global alignment of the two given networks using their maximum weight bipartite match with the scores as the weights. The GRAAL (GRAph ALigner) family (Kuchaiev and Pržulj, 2011) of global network alignment methods use the graphlet degree similarity to align two networks. Briefly, the graphlet-degree of a node counts the number of graphlets (i.e. induced subgraph) that this node touches, for all graphlets on 2 to 5 nodes. GRAAL (Kuchaiev *et al*., 2010) first selects a pair of nodes (one from each ofthe two given networks) with high graphlet degree signature similarity as the seed of the alignment, and greedily expands the alignment by iteratively including new pairs of similar nodes. H-GRAAL (Milenković *et al*., 2010), MI-GRAAL, andL-GRAAL algorithms alsobelong to the same family. The SPINAL algorithm (Aladaǧ and Erten, 2013) iteratively grows the alignment based on apriori computed node similarity score. MAGNA (Saraph and Milenković, 2014) optimizes the edge conservation between two networks using a genetic algorithm. There are several other methods for pairwise network alignment (Kelley *et al*., 2004; Phan and Sternberg, 2012; Gülsoy et al.,2012;Hasan and Kahveci, 2015;Patro and Kingsford, 2012; Neyshabur *et al*., 2013; Sun *et al*., 2015; Hu *et al*., 2013). Although the underlying algorithms of these methods vary, the end goal is similar to those discussed above.

Several algorithms address the multiple network alignment (Alkan and Erten, 2013; Ibragimov *et al*., 2014; Sahraeian and Yoon, 2013). IsoRankN (Liao *et al*., 2009) extends IsoRank. It adopts spectral clustering on the induced graph of pairwise alignment scores. The algorithm developed by Shih et. al. (Shih and Parthasarathy, 2012) is a seed-expansion heuristic that first selects a set of node pairs with high similarity scores using a clustering algorithm, and then expands these pairs by aligning nodes that maximizes the number of the total conserved edges of aligned nodes.

INQ (Hasan and Kahveci, 2014) aligns a dynamically evolving query network with one static target network. It uses ColT (Hasan and Kahveci, 2013) to find an initial alignment of the initial query, then it observes the differences between the topologies of the already aligned query network and the new query network, and finally, uses these differences to refine the alignment found for the previous query and generate alignment of the current query network. DynaMAGNA++ (Vijayan *et al*., 2017) aligns two dynamic networks. It assigns a value to each node based on how the incident edges and graphlets change through dynamic events. It assigns each node a value based on dynamic graphlet degree vector (DGDV) of graphlets up to size four. It considers a pair of nodes from two networks similar if their DGDVs are similar.

## 3 Problem Formulation

In this section, we develop anew scoring function, *score*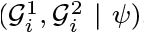, that integrates the similarities of the aligned nodes and their evolving topologies, and includes a penalty for each disconnected component in the aligned subnetworks of the target network at each time point. Next, we introduce the terminology and discuss how we drive our scoring function.

Given a network *G* = (*V, E*) and a subset of nodes 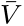, we define the induced subnetwork of 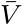 in *G* as the nodes in 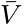 and all incident edges (i.e., 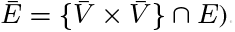. We denote this induced network as 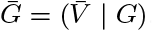. We say two nodes *u* and *v* in *G* are connected if there exists a path between *u* and *v* in *G*. We say a subset of nodes in *G* form a *connected component* if all pairs of nodes in that subset are connected in *G*. We define a subset of nodes 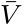 in *G* as a *maximum connected component* if the following conditions hold: (i) 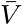 is a connected component in *G*, and (ii) there is no node in 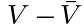 which is connected to a node in 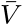. In the rest of the paper, we use the term “connected component” instead of “maximum connected component”. We denote the number of connected components of a given network *G* with *NCC*(*G*).

Given two temporal networks with *t* time points, 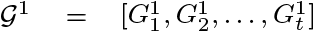 and 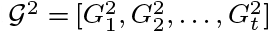, we denote the similarity between a pair of nodes *u* ∈ *V*^1^ and *v* ∈ *V*^2^ at time point *i* (1 ≤ *i* ≤ *t*) with *S_i_*(*u, v*).We use an existing pairwise alignmentmethod to calculate *S_i_*(*u, v*). The alignment function *ψ* maps all nodes in *V*^1^ to a subset of the nodes in *V*^2^. We denote this subset with Ψ(*V*^1^) (i.e. Ψ(*V*^1^) = {ψ(*u*)|∀*u* ∈ *V*^1^}). We note that *ψ* yields an induced subnetwork 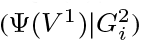 of 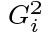 for each time point *i*, and each induced subnetwork 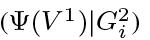 forms one or more connected components. See Figure 2(a) for an illustration of this latter point. We denote the number of connected components of the induced subnetwork 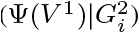 at time point *i* as 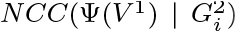. If the number of connected components at time point *i* is greater than one then the corresponding induced subnetwork is disconnected. We incur a penalty to account for the missing edges which would connect the disconnected components, and apply this penalty to each disconnected component.

**Fig. 2.**
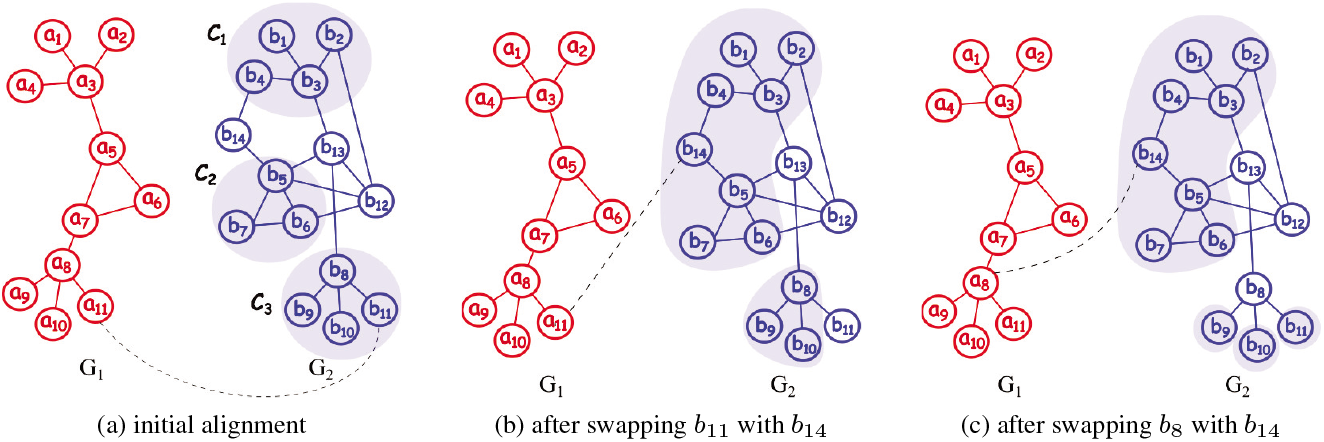
This figure represents an alignment between two networks *G*_1_ and *G*_2_. Each node in the query network *G*_1_ has a one-to-one mapping with a node in the network *G*_2_. The dashed line between two nodes emphasizes that they are mapped to each other. (a) This represents a hypothetical alignment where *a_i_* is aligned with *b_i_* for all 1 ≤ *i* ≤ 11. The induced subnetwork of the aligned nodes in *G*_2_ forms three connected components; *C*_1_ = {*b*_1_, *b*_2_, *b*_3_, *b*_4_},*C_2_* = {*b*_5_, *b*_6_, *b*_7_}, and *C_3_* = {*b*_8_, *b*_9_, *b*_10_, *b*_11_}. Gap nodes are {*b*_12_, *b*_13_, *b*_14_}. (b) After swapping *b*_11_ with *b*_14_. This swapping results in two connected components in *G*_2_. (c) After swapping *b*_8_ with *b*_14_. The aligned nodes in *G*_2_ form four connected components.

The minimum number of edges needed to join 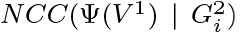 connected components is 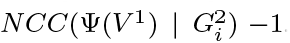. We penalize each edge insertion with a constant value denoted with *δ*, where *δ* ≥ *S_i_*(*u, v*), ∀ *u* ∈ *V*^1^, *v* ∈ *V*^2^ and *i* ∈ {1, 2, …, *t*}. We define the score of the alignment *ψ*() at time point *i* as: *score*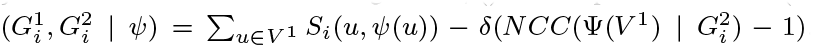. We define the temporal network alignment as

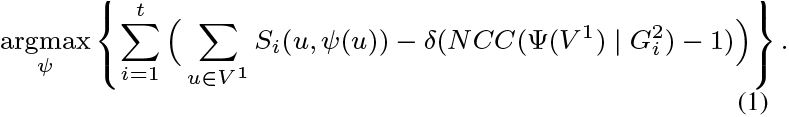

## 4 Methods

### Overview

Our algorithm for solving the temporal network alignment problem has two phases. The first phase finds an initial alignment between the input networks 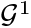 and 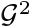 using the similarity score between pairs of aligned nodes across all time points. The induced subnetwork of 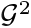 obtained by this alignment may be disconnected since this phase ignores the penalty incurred by edge insertions. The second phase reduces the number of connected components, improving the alignment score. In the second phase, we improve the alignment between the input networks by *swapping* a subset of the nodes in 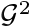 that are aligned with nodes in 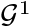 with other nodes in 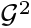. In order to swap a node *v_i_* ∈ Ψ(*V*^1^) with *v_j_* ∈ *V*^2^ – Ψ(*V*^1^), we update the alignment function *ψ*() to 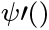 such that ∀ *u* ∈ 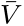 one of the two conditions is satisfied: (i) 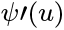 = *v_j_* if *ψ*(*u*) = *v_i_* and (ii) 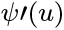 = *ψ*(*u*) if *ψ*(*u*) ≠ *v_i_*. Figure 2 illustrates this. Here, initially *b*_11_ is aligned to *a*_11_ (Figure 2(a)). Swapping *b*_11_ with *b*_14_ updates the alignment function so that *b*_14_ is aligned to *a*_11_ (Figure 2(b)). We observe that this swapping reduces the number of connected components in the induced subnetwork of *G*_2_ by one. Notice that if we swap *b*_8_ with *b*_14_ (instead of *b*_11_ with *b*_14_) then the number of connected components increases (Figure 2(c)).

We note that the number of connected components may simultaneously decrease at one time point and increase at other time points when we swap two nodes. We prove that the problem of finding the subset of node swaps that minimizes the number of connected components across all time points is NP-hard. We give areduction from the Maximum Coverage problem (Feige, 1998) to this problem later in this section.

### Algorithm details

Tempo takes two networks (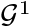 and 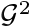) and the maximum number of allowed swaps (denoted as *k*) as input. In the following, we explain the two phases of our method in detail.

Phase I (Initialization). Here, we construct an initial alignment of 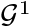 and 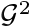. There exists several algorithms to perform pairwise alignment of two static networks at a single time point. Each of these methods assign similarity scores to all node pairs (one from the first network and one from the second) and then choose the alignment that maximizes the total score of all aligned node pairs. We adopt one of these methods to obtain the similarity scores of each network pairs (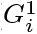, 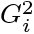) at each time point *i*, and use the outputted scores to calculate an initial alignment. We denote the similarity of the node pair (*u, v*), *u* ∈ *V*^1^ and *v* ∈ *V*^2^ generatedbysuch method at the *i*th time point with *S_i_*(*u, v*).

We generate an initial alignment *ψ*_0_ as follows. We first construct a weighted bipartite network *G_bp_* = (*V*^1^, *V*^2^, *ε*) as follows: we insert an edge in *G_bp_* between each pair of nodes (*u, v*) such that *u* ∈ *V*^1^ and *v* ∈ *V*^2^. We set the weight of the edge (*u, v*) as the similarity between nodes *u* and *v* aggregated over all time points. We denote the similarity as 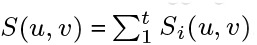. The maximum-weight bipartite matching algorithm maps each node in *V*^1^ to a node in *V*^2^ (Papadimitriou and Steiglitz, 1998). This mapping represents the initial alignment, *ψ*_0_. We call the nodes in *V*^2^ that are not mapped to any node in *V*^1^ as *gap nodes* and denote with *F* = *V*^2^ – Ψ(*V*^1^).

Phase II (select *k* swapping pairs). Here, we describe our dynamic programming algorithm that selects a set of *k* swaps which maximize the alignment score by reducing the number of connected components in the induced alignment across all time points of 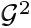 (see Equation 1).

We denote a set of *r* swaps with Δ = {(*u*_1_, *v*_1_), (*u_2_, v*_2_),…, (*u_r_, v_r_*)} with ∀*i* ≠ *j, u_i_* ≠ *u_j_* and *v_i_* ≠ *v_j_*. We denote the alignment after applying the swaps in a given set Δ as *ψ*_Δ_ Let us denote the optimal set of *r* swaps for the alignment *ψ* with *solution*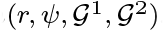 Also, for a given *u_i_* ∈ Ψ(*V*^1^), we denote the optimal set of *r* swaps for the alignment *ψ* which contains the swap pair (*u_i_, v_i_*), ∃*v_i_* ∈ *F*, with *solution*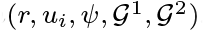

Our algorithm works iteratively. In the first iteration, our algorithm selects one swapping pair for each aligned node *u_i_* ∈ Ψ(*V*^1^) as

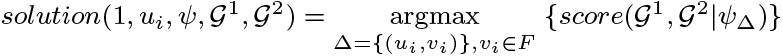

At each subsequent iteration *r* where 2 ≤ *r* ≤ *k*, for each aligned node *u_i_* ∈ Ψ(*V*^1^), our algorithm selects a set of *r* swapping pairs denoted with *solution*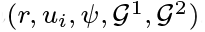 by adding one swapping pair (*u_i_, v_i_*), ∃*v_i_* ∈ *F*, to the previously selected *r* – 1 pairs as follows.

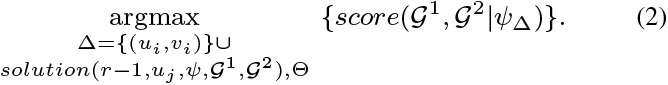

Here Θ represents the necessary conditions to include the (*u_i_, v_i_*) swap pair with a set of *r* – 1 swap pairs as

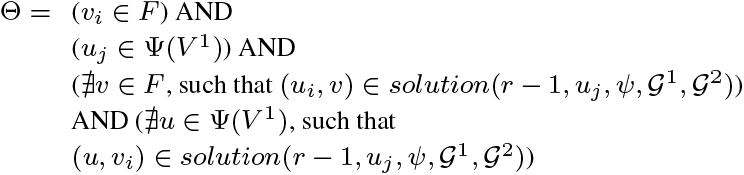

The first condition above ensures that node *u_i_* is swapped with a gap node and the second ensures the dynamic programming iterates over all size *r* – 1 swap sets for all aligned nodes of 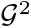. The third condition ensures that the aligned node *u_i_* has not already been swapped in the *r* – 1 sized swap set. The final condition is the dual of the previous one, as it ensures that the gap node *v_i_* has not already been swapped in the *r* – 1 sized swap set. When these conditions hold, the two nodes *u_i_* and *v_i_* can be swapped and included into the existing set of *r* – 1 swaps without conflicting with any of the existing swaps.

We report the output of the algorithm at end of the *k*th iteration as set of *k* swaps with the highest alignment score using equation

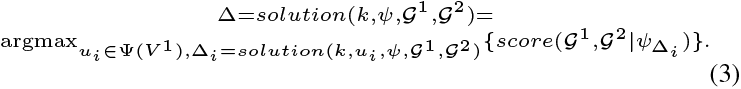

We represent the set cardinalities |*V*^1^|, |*V*^2^|, and |*F*| with *m, n, l*, respectively. The complexity of our algorithm is 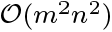 + 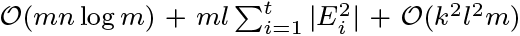. We provide the derivation of this complexity in Section A.1 of the appendix. We note that *k* ≤ 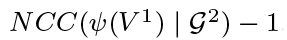. This value is either given as input or we set it to 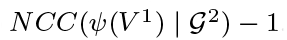.

### Proof of correctness

Here, we formally proof the correctness of our algorithm. We say that swapping the pair of nodes (*u_i_, v_i_*) is *proper* if that the swapping does not increase the number of connected components of the aligned nodes. We first prove that our algorithm will always find a proper swapping node *u_i_* from the set of aligned node for each gap node *v_i_*. We first present a lemma which is necessary for the proof of our first theorem. Let us denote the degree of a node *v* (i.e. number of edges connected to this node) within a component *C_i_* = (*V_c_, E_c_*) of the induced subnetwork 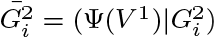 at time point *i* with the function *deg(v|C_i_)*.

Lemma 1. *Given an undirected subnetwork of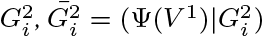 where |*V_c_*| = *z* and 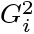 is acyclic network (has no cycle) within its topology, then Σ_*v∈C_i_*_ *deg*(*v*|*C_i_*) = 2(*z* – 1)*

*Proof*. Since *C_i_* is a connected subnetwork with no cycles, the number of edges in *C_i_* equals *z* – 1 edges. Each edge belongs to an undirected network increases the sum of the network nodes degrees by two. Thus, Σ_*v∈C_i_*_ *deg*(*v*|*C_i_*) = 2(*z* – 1)

Lemma 2. *Given a gap node *v_i_* that connects at least two connected components, there exist at least one aligned node *u_i_* which we can swap with *v_i_* without increasing the number of connected component*.

*Proof*. We formally prove this by induction on the size of connected components that *u_i_* belongs to.

Base case. We consider a component *C_i_* = (*V_c_, E_c_*) where |*V_c_*| =2 and *v_i_* is connected to *C_i_* through *u_j_*, and assume *u_i_* belongs *C_i_*. If we swap *v_i_* with *u_i_*, then *C_i_* will contain *u_j_* and *v_i_* which corresponds to one component. Thus, the number of connected components of *C_i_* is still one after swapping.

Induction hypothesis. We assume there exists a node *u_i_* for all components of size *q* nodes that can be swapped without disconnecting its component. We consider two cases of one component *C_i_* where *v_i_* is connected to through *u_j_*. The first case is when *C_i_* contains at least one cycle with the set of nodes, *V_c1_* = {*v*_1_, *v*_2_,…, *V_n_*}. It follows that for each node *u_i_* ∈ *V_c1_* and *u_i_* ≠ *u_j_, u_i_* can be swapped with *v_i_* without disconnecting *C_i_*. In the second case, *C_i_* represents acyclic network with no cycles. Next, we prove our theorem in this case by contradiction. First, we assume that the number of nodes in *C_i_* with degree equal to 1 is less than 2. Consequently, Σ_*v∈C_i_*_ *deg*(*v*|*C_i_*) ≥ 2(*z* – 1), which contradicts Lemma 1. Thus, the number of nodes in *C_i_* with degree equal to 1 is at least 2 nodes and thus, ∃*v,w ∈ C* st. *deg*(*v*|*C*) = 1 and *deg*(*w*|*C*) = 1 and *v* ≠ *w*. Therefore, we can swap *v_i_* with either *v* or *w*.

Next, we prove that swapping a gap node *v_i_* with an aligned node *u_i_* at each iteration will increase the alignment score *score*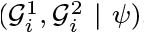, showing that the alignment score will always improve by our dynamic programming algorithm.

#### Theorem 1.

Given a value of *δ* where *δ* is greater than or equal to *S*(*ψ*(*u_i_*), *u_i_*) for all *u_i_* ∈ *V*^2^. At each iteration of our algorithm, *score*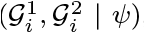 monotonically increases.

#### Proof

We assume that our algorithm chooses one pair of nodes to swap; a gap node *v_i_* and aligned node *u_i_* which will connect *x* number of components. We note that the condition *x* ≥ 2 must be satisfied for *v_i_* to be considered for swapping. Also, it follows from Lemma 2 that if we swap *v_i_* and *u_i_* then the number of connected components will not increase. Thus, the difference in the score equals *D* = *δ*(*x* – 1) – *p_uv_* where *p_uv_* is the difference in pairwise score from swapping (i.e. *p_uv_* = *S*(*ψ*(*u_i_*), *u_i_*) – *S*(*ψ*(*u_i_*), *v_i_*)). Since *δ* is greater than or equal to *S*(*u, v*) ∀ *u* ∈ *V*^1^ and ∈ *V*^2^, then *δ*(*x* – 1) ≥ *p_uv_*. Consequently, *D* ≥ 0 and 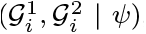 will not decrease.

### ProofofNP-hardness

Here, we prove that our problem is NP-hard. To do that, we reduce the *Maximum Coverage Problem (MCP)*, which is known to be NP-hard (Karp, 1972), to our problem. Given a positive integer 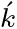 and a collection of sets, *S* = {*S*_1_, *S*_2_,…, *S_m_*}, MCP seeks the subset *Ś* 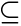 *S* such that |*Ś*| ≤ 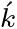 and the number of covered elements |⋃_*S_i_∈Ś*_ *S_i_*| is maximized.

We reduce MCP to an instance of our problem. Let *U* = {*x*_1_, *x*_2_, …, *x_n_*} be the union of elements in *S* (i.e. *U* = |⋃_*S_i_∈Ś*_ *S_i_*|). We construct a target temporal network 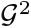 with one time point *G*^2^ = (*V*^2^, *E*^2^) as follows. We initialize *G*^2^ as *V*^2^ = 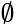 and *E*^2^ = 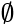. Next, we add a node *a_j_* in *G*^2^ for each element *x_j_* ∈ *U*. Also, for each set *S_i_* ∈ *S*, we add two nodes *f_i_* and *b_i_* in *V*^2^. Formally, *V*^2^ = {*a*_1_, *a*_2_,…, *a_n_*}⋃{*b*_1_, *b*_2_,…, *b_m_*}⋃{*f*_1_, *f*_2_,…, *f_m_*}.Next, we populate the set of edges *E*^2^. To do that, for all *S_i_* ∈ *S* and *x_j_* ∈ *S_i_*, we insert the edge (*f_i_, a_j_*) in *E*^2^. In addition, for all pair of sets *S_i_, S_j_* ∈ *S*, where *i* < *j*, we insert the edge (*f_i_, f_j_*) in *E*^2^. Finally, for a given query network *G*^1^ = (*V*^1^, *E*^1^), we construct the set of nodes in *G*^2^ aligned to those in *G*^1^ as Ψ(*V*^1^) = {*a*_1_, *a*_2_,…, *a_n_*}⋃{*b*_1_, *b*_2_,…, *b_m_*}. Thus, the set of gap nodes is {*f*_1_, *f*_2_,…, *f_m_*}. Notice that, the subnetwork of *G*^2^ induced by Ψ(*V*^1^) has *m*+*n* nodes but it contains no edges as all the edges in *G*^2^ are connected to a gap node by our construction. Thus, the alignment yields *n* + *m* connected components as each node in Ψ(*V*^1^) represents a component.

Recall that each swapping operation swaps an aligned node with a gap node. Also, recall that the optimization problem we solve for aligning temporal networks aims to find at most *k* swaps, such that after applying those swaps, the number of connected components *NCC*(Ψ(*V*^1^) | *G*^2^) is minimized (see Section 3). We call this optimization problem *minimum Connected Component Problem (mCCP)* in the rest of this proof. Next, we prove that MCP is maximized if and only if mCCP is minimized.

First, we prove that if there exists a solution to mCCP, then there exists a solution to MCP. In other words, we prove that minimizing mCCP maximizes MCP. Let us denote the nodes corresponding to the elements in a set *S_i_* with *A_i_* = U*_x_j_∈S_i__* {*a_j_*}. In our problem instance, a swap operation swaps *f_i_* with a node in the set *V*^2^ – *A_i_* – {*f_i_*}. This is because all nodes in *A_i_* are connected to *f_i_*, and thus swapping *f_i_* with a node not in *A_i_* ensures that all nodes in *S_i_* ⋃ {*f_i_*} form one connected component. Therefore, to minimize the number of connected components, we swap *f_i_* with one of the nodes which is not a part of this connected component. To ensure that, we swap *f_i_* with a node in the set {*b*_1_, *b*_2_, …, *b_m_*}. Since all nodes in this set are disconnected, swapping *f_i_* with any node in this set will yield the same number of connected components. Let us assume that the solution to mCCP performs *k* swaps. Following from the discussion above, without losing generality, we assume that these swaps are {(*f*_1_, *b*_1_), (*f*_2_, *b*_2_),…, (*f_k_, b_k_*)}. Notice that after these swaps, the nodes in 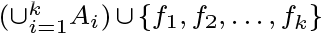 forms one connected component, and all remaining nodes are isolated. Let us denote the number of connected components after these swaps with *β*. Let us denote the number of nodes in 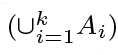 with *τ*. Notice that *τ* also reflects the number of elements covered in 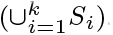. We have *β* = (*m* – *k*) + (*n* – *τ*) + 1.

In the formulation above, the first term (*m* – *k*) is the number of nodes *b_j_* which are not swapped with a gap node. Since all those nodes are isolated, each one forms a connected component by itself. The second term (*n* – *τ*) is the number of nodes *a_j_* which are not included in the set 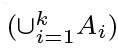. These nodes remain isolated even after swapping of nodes. The last term (i.e., 1) is the connected component containing the nodes in 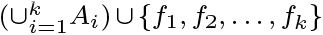. After minor algebraic manipulation, we rewrite the equation above as *β* = (*m* + *n* – *k* +1) – *τ*. In this equation, the parameters *m, n*, and *k* are input to the given mCCP problem, and thus we denote the first term above with the constant *c* = *m*+ *n*– *k*+ 1. Therefore,we have *β* = *c*–*τ*. In this equality the smaller the value of *β* is, the larger *τ* gets. Thus, minimizing the number of connected components *β* in mCCP maximizes the nuumber of elements covered in MCP.

Second, we prove that if there exists a solution to MCP, then there exists a solution to mCCP. In other words, we prove that maximizing MCP minimizes mCCP. Let us assume that the solution to MCP is *Ś* = 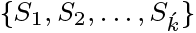. The number of elements coveredby this solution is 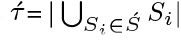. By constructing an instance of mCCP as described above, we have 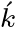 swaps denoted with the set 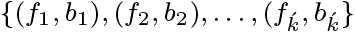. Consequently, after performing these swaps, the nodes in 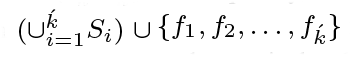 forms one connected component, and all the remaning nodes are isolated. Let us denote the number of connected components with 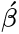. We have 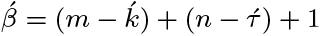.

After minor algebraic manipulation, we rewrite the equation above as 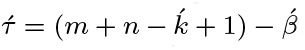. Since *m, n*, and 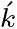 are input parameters, we have 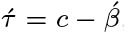, where *c* is a constant 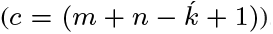. In this equality, the larger the value of 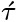 is, the smaller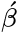 gets. Thus, maximizing 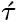 in MCP results in maximizing 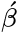 in mCCP.

Lastly, the proof we describe above reduces an instance of MCP to an instance of mCCP in polynomial time and space as it requires only building a network with 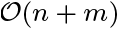 nodes and edges. Thus, we conclude that the mCCP problem is NP-hard.

## 5 Results and Discussion

We evaluate the performance of our algorithm on synthetic and real data. Next, we describe both datasets in detail.

**Real Dataset**. We obtain our real dataset from two sources. The first one is the human brain aging dataset (Berchtold *et al*., 2008). Recall that this dataset contains gene expressions of 173 samples obtained from 55 individuals spanning 37 ages from 20 to 99 years. The ages in this dataset are not uniformly spaced. In order to bring consecutive time gaps to a more uniform values, we remove two data points which have an age gap of more than 5 years from their successive age values, leading to 35 ages. We select five temporal networks each having seven time points. Next, we explain how we do that for the first temporal network. We start with the first (i.e., youngest) time point in the aging data. We then skip the next four time points and take the sixth time point in aging data iteratively until we have seven time points. Similarly, for 1 < *j* ≤ 5, we select the *j*th temporal network starting from the jth time point. In this manner, we form five nonoverlapping and interleaved temporal networks. In order to integrate static PPI network with gene expression data to form age-specific PPI networks, we set a cut-off on the gene-expression value. All the interactions that have a lower transcription value for either or both the proteins are removed from the corresponding age-specific network. We use the protein-protein interaction (PPI) network data from BioGRID (Breitkreutz *et al*., 2007). For the second source, we select phenotype specific query temporal networks from this dataset. We use two neurodegenrative disorders which are conjectured to be age-related (Alzheimer’s and Huntington’s) and a third one which we expect to be less prone to aging (Type II diabetes). We retrieve the gene sets specific to these three diseases from KEGG database (Ogata *et al*., 1999). We form three query PPI temporal networks by keeping only the interactions where both the interactors are from each of the three phenotype-specific (Alzheimer’s, Huntington’s or Type II Diabetes) gene set.

**Synthetic dataset**. We generate synthetic networks to observe the performance of our method under a wide spectrum of parameters classified under two categories; (i) network size and (ii) temporal model parameters, namely number of time points, temporal rate, and cold rate. We vary the targetnetworksize to takevalues from {100, 250, 500, 750,1000}.Wefix the network density to two edges per node on the average (i.e., mean node degree is set to four). We randomly select 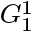 as a connected subnetwork of 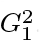. We set the size of the query network to 50 nodes. We generate target network 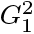 using Barabási-Albert (BA) (Barabási and Albert, 1999) model as this model produces scale-free networks. In order to explain the parameters in the second category, we describe how we generate the query and target networks 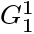 and 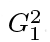 at the first time point. We then explain how we use the parameters in this category to build the query and target networks at the remaining time points.

We generate the subsequent networks for the remaining time points using the three parameters in the second category above as follows. The first parameter is the number of time points *t* in 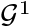 and 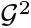. We use 5, 10, 15, and 20 time points in our experiments. Recall that we select a subnetwork of the target network 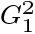 as the first query network 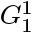. We mark all nodes and edges in 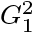 within this subnetwork as *cold* nodes and edges respectively. We mark all other nodes and edges in 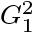 as *hot*. Next, we iteratively generate the networks 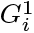 and 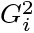 at the *i*th time point (*i* > 1) from 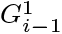 and 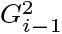 respectively as follows. Letus denote temporal and cold rates (two real numbers) with *∈* and *∈^c^* respectively such that 0 ≤ *∈^c^* ≤ *∈* ≤ 1. Let us denote the ratio of cold edges to the total number of edges in the target network 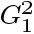 with *γ*. We calculate the hot rate, denoted with *∈^h^*, from temporal rate and cold rate as *∈^h^* = (*∈*– *∈^c^γ*)/ (1–*γ*). Conceptually, hot and cold rates model the rate of evolution of hot and cold edges between two consecutive time points respectively. More specifically, for each subsequent time point *i*, we generate 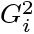 by randomizing 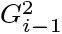 as follows. We iterate over all edges in 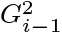. For each edge *e*, if it is a cold edge we remove it with probability *∈^c^* and insert a new edge between two randomly chosen cold nodes. If *e* is a hot edge, we remove it with probability *∈^h^* and insert anew hot edge between two random nodes (with at least one being a hot node). We generate query networks at subsequent time points using almost the same procedure with the only difference being that all edges are cold. We generate datasets by varying *∈* and *∈^c^* to take the values {0.05, 0.1, 0.2, 0.4, 0.8} and {0.05, 0.1, 0.2} respectively. Foreach parameter setting we generate 10 target and query temporal networks.

Recall that, we generate the scoring matrix based on both homology and topology similarities. We generate the homology score between two pair of nodes *u* ∈ *V*^1^ and *v* ∈ *V*^2^ as follows. If *v* was originally selected as cold node and *u* is the same as v, then we generate a homology score between *u* and *v* from log-normal distribution (Johnson *et al*., 1994) with mean 2*μ* and standard deviation *σ*. Otherwise, we randomly generate the homology score between *u* and *v* from log-normal distribution with mean *μ* and standard deviation *σ*. In this way, we allow nodes in query network to be likely to align to nodes in the target network that were originally extracted from. In this paper, we set *μ* and *σ* to be 2 and 0.25 respectively. Notice that the homology scores do not change through time points, although topology scores do. Thus, evolution through time points of query and target networks may affect how the query is aligned to the cold region in the target network. We set the edge insertion penalty *δ* to be 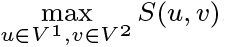

We compare the accuracy and running time of our algorithm against IsoRank, MAGNA++ and GHOST (see Appendix, Section A.3 for extended results). IsoRank, MAGNA++ and GHOST are designed to align two networks at a single time point. We therefore find the alignment using each of these methods at each time point, impose the alignment to all the time points and report the average. We analyze the biological significance of our results on real data by performing gene ontology analysis and exploring publication evidence. We implemented Tempo in C++, performed all experiments on a computer equipped with AMD FX(tm)-8320 Eight-core Processor 1.4 GHz CPU, 32 GB of RAM running Linux operating system, and used *α* = 0.7 unless otherwise stated.

### 5.1 Evaluation of recovered region

In this experiment, we compare the accuracyofthe alignment generatedby Tempo against that of IsoRank, MAGNA++, and GHOST. We recall that we select the original query network from a subset of nodes and their edges from the target network, and then evolve the query through time points. Here, we evaluate the accuracy by calculating the percentage of the aligned nodes from query network that are paired with the same nodes of the target network that they were originally selected from. We refer to this percentage as *recovered region*. We illustrate the results in Figure 3, which demonstrate that Tempo recovers high percentage of the query networks compared to other methods. As the temporal rate increases, the accuracy of Tempo improves dramatically while that of IsoRank remains nearly stagnant and while MAGNA++ and GHOST continue to generate alignments with low recovery rates. Growing the temporal rate while keeping the cold rate unchanged means that the topology of the query network (i.e., cold edges) is evolving slower than the rest of the temporal network (i.e., hot edges). This implies that Tempo can capture the variation in such evolutionary rate while competing alignment strategies which fail to do so.

**Fig. 3.**
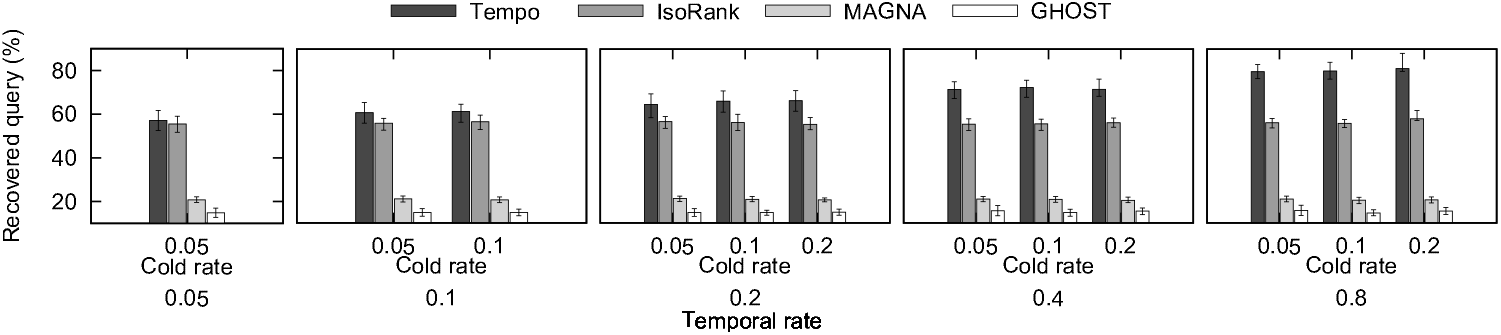
The percentage of recovered query in the resulting alignment varying *∈* and *∈^c^* to take the values {0.05, 0.1, 0.2, 0.4, 0.8} and {0.05, 0.1, 0.2} respectively. The x-axis shows temporal rate, *∈* and cold rate,*∈^c^* (these are the parameters used for constructing synthetic temporal network, with varying evolution rates. The y-axis shows the percentage of recovered query of IsoRank, MAGNA++, and GHOST against Tempo. The error bars show the 80-percentile of the recovered query based on the 10 repetitions of each parameters setting.

### 5.2 Evaluation of induced conserved structure

Next, we evaluate the topological quality of the alignment generated by Tempo through comparison with IsoRank, MAGNA++, and GHOST. For this purpose, we measure the shared topological structure between 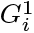 and 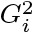 which is preserved under the alignment function *ψ* through all time points *i*. Induced conserved structure (ICS) measures the percentage of edges from 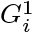 that are aligned to edges in 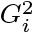 to the total edges of the induced subnetwork 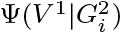, and is one of the most common measures of topological quality (Vijayan *et al*., 2015). Formally, 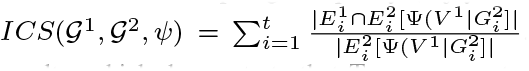. Figure 4 presents the results, which demonstrate that Tempo generates alignments with high quality based on ICS compared to other algorithms. We note that GHOST was created to optimize ICS, however, Tempo outperforms GHOST on this measure—especially when the temporal rate is high since the performance of GHOST degrades. We also evaluate our algorithm against other algorithms using the edge correctness (EC) measure (see Section A.3).

**Fig. 4.**
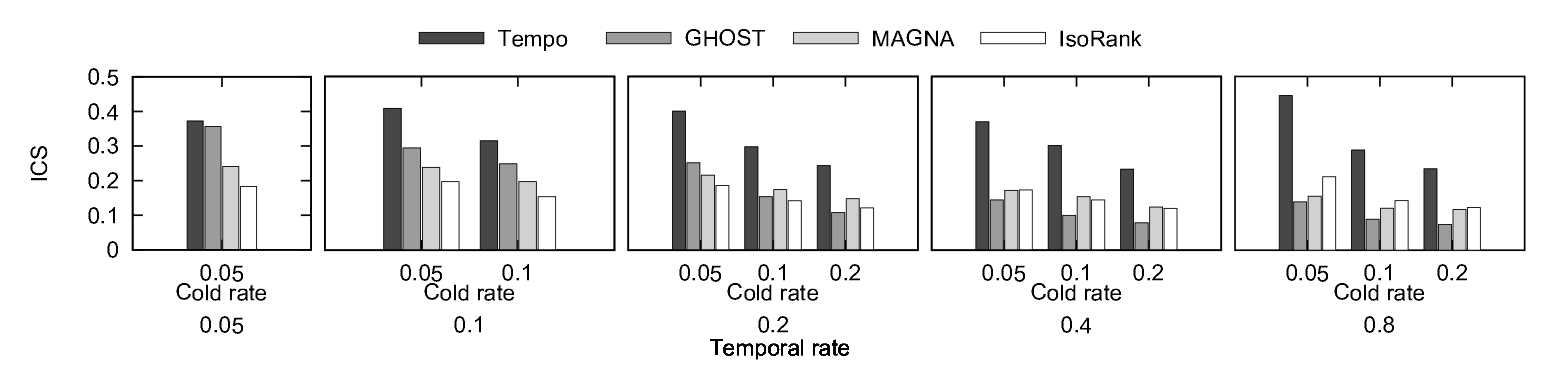
The induced conserved structure (ICS) score of the resulting alignment varying *∈* and *∈^c^* to take the values {0.05, 0.1, 0.2, 0.4, 0.8} and {0.05, 0.1, 0.2} respectively. The x-axis shows temporal rate, *∈* and cold rate, *∈^c^*. The y-axis shows the ICS score of GHOST, MAGNA++, and IsoRank against our method (Tempo).

### 5.3 Evaluation of statistical significance of the alignment

We compare the statistical significance of the alignments generated by Tempo against that of existing methods. In order to ensure that our experiments do not give any advantage to our algorithm, we use IsoRank to generate initial alignments for Tempo and thus, compare the statistical significance against IsoRank only.

**Varying evolution rate**. In this experiment, we evaluate the effect of varying the temporal rate (*∈*) and cold rate (*∈^c^*) on the significance of the score of the alignments produced by Tempo and that of IsoRank. We generate synthetic networks of sizes {100, 250, 500, 750, 1000} and 20 time points. We fix the network density to two edges per node on the average, and vary *∈* and *∈^c^* (*∈* ≤ *∈*) to take the values {0.05, 0.1, 0.2, 0.4, 0.8} and {0.05, 0.1, 0.2}, respectively. Next, we randomly selected 50 nodes from target network 1,000 times, and calculate the alignment score of each, i.e., each random selection corresponds to an alignment. We calculate the mean and standard deviation of these 1,000 scores and generate the z-score of the alignment generatedby Tempo using this mean and standard deviation. Hence, we denote the score generated from our methodby *S*\*, and denote the mean and standard deviation of 1,000 scores generated from the random selections with *S_μ_* and *σ*, respectively. We calculate the z-score of our method as (*S*\* – *S_μ_*)/*σ*. We calculate the z-score of the IsoRank method in a similar manner. Figure 5 presents the average z-score values across all target network sizes. The results show that as we increase the temporal rate, the z-score of Tempo significantly increases while the z-score of IsoRank increases by small amount. As the evolution rate increases, the topology of the alignment found by Tempo differs significantly from the topology of rest of the network, and thus, it becomes more challenging to find the correct alignment. However, Tempo continues to generate accurate and significant results especially for large evolution rates unlike IsoRank which considers each single time point independently. We observe the same pattern as we increase cold rate. **Varying time points**. In this experiment, we evaluate how the z-scores of Tempo and IsoRank differ as the input networks evolve and deviate from each other. More specifically, we consider aligning the query network with each of the four target sets we have which have evolving time points (i.e. older ages) as we move to later target sets. First, we measure the z-score of aligning thequery to the firsttargetset(i.e., containing timepoints 2, 7, 12, …) then we measure the z-score of aligning the query to the second target set (i.e., containing time points 3, 8, 13, …) and so on. We present the average z-score across all temporal and cold rates. Figure 6(a) presents the results. The results show that Tempo continues to generate alignment with high score significance as we evolve the network. We observe the same pattern for IsoRank, however, Tempo outperforms IsoRank—especially when the time points are distant. This confirms the fact that as the target and query networks evolve and deviate from each other, Tempo is able to take into account the evolution through consecutive time points and generate accurate alignments that persist.

**Fig. 5.**
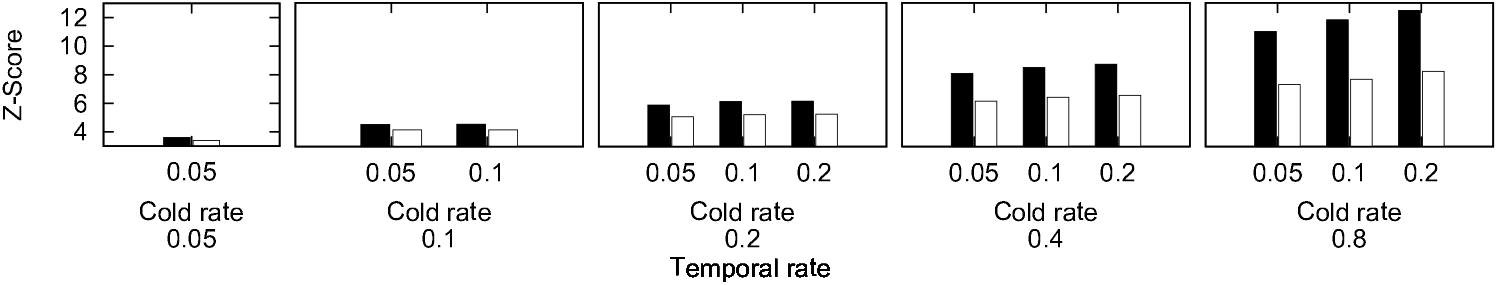
The average z-score of Tempo across network sizes {100, 250, 500, 750, 1000} varying *∈* and *∈_c_* to take the values {0.05, 0.1, 0.2, 0.4, 0.8} and {0.05, 0.1, 0.2} respectively. The x-axis shows temporal rate, *∈* and cold rate, *∈_c_*. The y-axis shows the z-score of IsoRank (white) against Tempo (black).

**Fig. 6.**
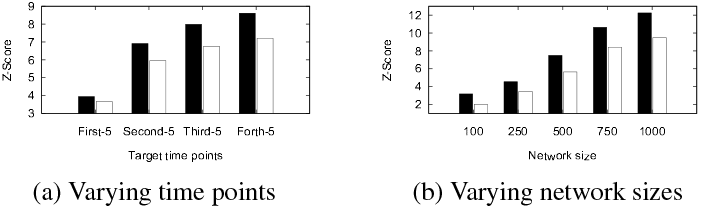
The average z-score of Tempo (black) against IsoRank (white) (a) varying target time points, the x-axis shows time point selected, and (b) varying network size, the x-axis shows network sizes in terms of number of nodes.

**Varying network size**. In this experiment, we compare the significance of the alignment generated by Tempo against IsoRank as the target network size increases and the query becomes small with respect to the target. We average the z-score across all evolution rates and vary target network size to take values {100, 250, 500, 750, 1000}. Figure 6(b) presents the results, which show that the significance of the alignment (best alignment) increases as we increase the size of the underlying target network. We expect this behavior since we compare the aligned nodes (50 nodes) to a random selection of 50 nodes from the underlying target network. Thus, the chance of selecting the best alignment decreases. That said, Tempo was able to identify the accurate alignment which results in high significant values.

### 5.4 Evaluation on real data

Next, we evaluate Tempo on the real data. We first evaluate the significance of alignment score using Tempo. We calculate the z-score by comparing the score of aligned nodes to the score of 1,000 randomly selected alignments of the same number of nodes. We compare our results to those of IsoRank.

We repeat this experiment for three different disease network queries: Alzheimer’s, Huntigton’s and Type-II diabetes. Figure 7 shows the results. Our results demonstrate that Tempo yields highly significant alignments, and outperforms IsoRank in terms of z-score. We also observe that z-scores of non-age related disease (diabetes) is lower than those of age-related diseases (i.e. Alzheimer and Huntington’s). Although there are some fluctuations in the z-score with growing time gap between query and target networks, we observe that the z-score tends to increase for Alzheimer’s and Huntington’s disease unlike the Type-II diabetes. This suggests that age-related pathways have higher evolution rate than other pathways. Thus, we conjecture that Tempo, which takes all time points into consideration, is suitable for capturing evolving topologies.

**Fig. 7.**
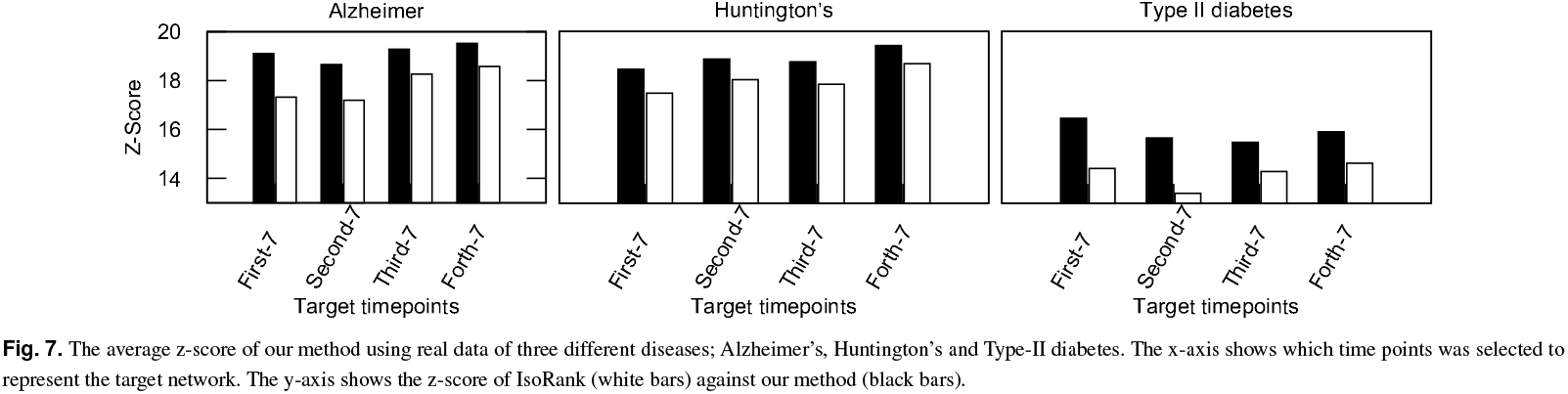
The average z-score of our method using real data of three different diseases; Alzheimer’s, Huntington’s and Type-II diabetes. The x-axis shows which time points was selected to represent the target network. The y-axis shows the z-score of IsoRank (white bars) against our method (black bars).

Next, we consider the biological significance of our results by identifying aligned gene pairs in which the aligned genes are different, and determining prior evidence that these gene pairs are biologically relevant. We use Tempo to identify 4, 4 and 6 such pairs for Alzheimer’s, Huntington’s and Type-II diabetes, respectively. We note that Alzheimer’s, Huntington’s and Type-II diabetes query sizes are 39, 36, and 23. Thus, thepercentagesofthe different genes found to all the genes in the alignment are 10% to 26%. IsoRank only mapped genes to themselves, suggesting that IsoRank only considers static topologies while our algorithm could map genes based on homological similarities as well as evolving topologies. MAGNA++ and GHOST could only map few genes to themselves while other mapped genes were poorly related (see appendix, Section A.3).

For each combination of disease and differently mapped gene pairs identified by Tempo, we first search PubMed for publication evidence specific to that disease. For instance, in case of Alzheimer’s disease, the gene DAB1 that was selected by Tempo and was identified as a potential gene that encode proteins related to functions in biological pathways relevant to the disease (Gao *et al*., 2015). Genes found by Tempo for type II diabetes, for example gene ACTA1, has remarkable change in gene expression value that was observed for the in diabetic samples compared to non-diabetic samples (Nashida *et al*., 2013). Moreover, significant up-regulation of GRB2 is observed in transgenic samples compared to controls (Burdon *et al*., 2015).

In order to determine the biological processes of the aligned genes found by Tempo in gene aging dataset, we perform the gene ontology analysis of the aligned genes in target network using Gene Ontology Consortium (Consortium *et al*., 2004). We identify the biological processes or signaling pathways that play significant roles in the disorder. We calculate how many related pathways found by our method (Tempo) against MAGNA and GHOST and their significance. We also counted the frequency of those pathways when used different range of time points. Table 1 present the results. We find references of certain pathways that are related to specific neurodegenerative disorders (Alzheimer’s and Huntigton’s diseases). For genes we identify when we use Alzheimer’s disease as a query network, we find two pathways, namely *Alzheimer disease-amyloid secretase* and *Alzheimer disease-presenilin* are related to Alzheimer’s disease (Mattson, 2004). Various growth factors alter the brain development process at younger age, that manifest as a variety of risk factors at an older age and eventually results in aging-related diseases such as Alzheimer’s and Huntigton’s diseases (Bartzokis, 2004). For the genes we identify when we use type II diabetes phenotype as a query, we find two pathways that they are commonly associated with type II diabetes (Liu *et al*., 2007) namely *Insulin/IGF pathway-protein kinase B signaling cascade and Insulin/IGF pathway-mitogen activated protein kinase kinase/MAP kinase cascade*. On the other hand, MAGNA or GHOST found at most one pathway with very low significance and did not appear through all tested target networks (see Table 1). In conclusion, studying temporal networks in general and human aging specifically using Tempo enables us to identify age related genes from non age related genes successfully. More importantly, Tempo takes the network alignment problem one huge step forward by moving beyond the classical static network models.

**Table 1.**
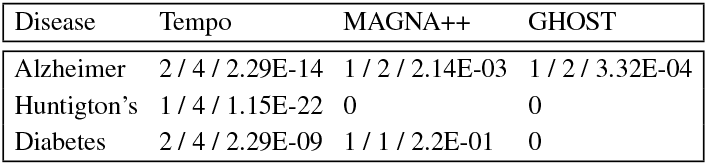
Number and significance of functional pathways associated with the underlying disease observed among the aligned genes of target network. Each cell lists the results in the form x/y/z. Here, x represents number of pathways identified, y denotes the number of time points at which these pathways are observed, and z is the statistical significance (p-value) of the least significant of these pathways. The cell with the value 0 implies that no pathways were found.

